# ECHOS enables spatial epigenome profiling at subcellular resolution

**DOI:** 10.64898/2026.03.26.714421

**Authors:** Qiqi Cao, Qianlan Xu, Yusuke Ueda, Shreya Rajachandran, Manjita Sharma, Xin Zhang, Mala Mahendroo, Edward J. Grow, Haiqi Chen

**Affiliations:** Cecil H. and Ida Green Center for Reproductive Biology Sciences, University of Texas Southwestern Medical Center, Dallas, TX, USA; Department of Obstetrics and Gynecology, University of Texas Southwestern Medical Center, Dallas, TX, USA

## Abstract

Biological structures and the epigenome are intertwined. For example, complex tissues are often the combined products of various groups of spatially patterned cell types with distinct epigenetic states. Furthermore, chromatin at various subnuclear locations within a cell often differ in their epigenetic properties. Thus, a systematic understanding of the relationship between the epigenome and its spatial distribution across biological scales would inform tissue and cellular functions as well as gene regulatory mechanisms. Yet, spatially resolved epigenome profiling—particularly at subcellular resolution—remains technically challenging. Here, we present <u>E</u>pigenetic <u>C</u>UT&<u>T</u>ag via <u>H</u>igh-resolution <u>O</u>ptical <u>S</u>election (ECHOS), a platform that combines high-resolution imaging and high-throughput sequencing to enable precise, spatially targeted epigenetic profiling across biological scales. At the cellular scale, ECHOS generates high-quality DNA-binding protein and histone modification datasets that show strong concordances with datasets from ChIP-seq and CUT&Tag experiments. Further optimization of ECHOS (ECHOS+) enables the characterization of the histone modification landscape of chromatin at the sub-micron resolution. Using ECHOS+, we revealed distinct gene regulatory logics at different layers of human ectocervical epithelium. We also showed that micronuclei—small nucleus-like structures formed by mitotic errors—exhibited a different epigenetic state from the same chromosome regions on the intact nuclei. Finally, we found that human aging altered the epigenetic state of the inactive X chromosome located in a subcellular nuclear structure called the Barr body, which may contribute to genes escaping X chromosome inactivation during female aging. Together, ECHOS and ECHOS+ represent a scalable and generalizable framework for spatial epigenomic analyses, with broad potential applications in various domains of biology.

## Introduction

The spatial distribution of the epigenome has been increasingly appreciated as an important gateway into understanding tissue and cellular functions^1,2^. For example, at the tissue level, layer-specific H3K4me3 profiles have been identified at the promoter regions of certain genes within the adult mouse brain cortex^3^. At the subcellular level, the gene-rich, transcriptionally active chromatin is more centrally located within the nucleus, whereas the gene-poor, transcriptionally repressed chromatin preferentially occupy the nuclear periphery and are enriched for repressive histone modifications^4^. Thus, being able to characterize the epigenetic states of chromatin in a spatially resolved manner would allow a deeper understanding of tissue and cellular functions.

Despite its biological importance, spatial epigenome organization especially at the subcellular level has remained largely inaccessible to experimental measurements. Bulk epigenomic assays such as ChIP-seq^5^, CUT&RUN^6^ and CUT&Tag^7^ require pooled materials and therefore lose anatomical and nuclear context. Single-cell epigenomic profiling methods reveal cell-level heterogeneity but also fail to retain spatial information due to tissue disassociation^8–13^. Physical microdissection of cells (e.g., laser-capture microdissection) provides limited spatial resolution, suffers from sample loss and contamination, and is not applicable to probing subcellular structures^14,15^.

Emerging technologies have begun to address the need for spatially resolved epigenome profiling. Microfluidic barcoding approaches such as Spatial-CUT&Tag enables spatial epigenomic measurements by patterned delivery of molecular tags but require specialized microfabricated devices and have limited spatial resolution (20 μm)^16^. Multiplexed fluorescence in situ hybridization–based approaches, such as epigenomic MERFISH, can report the spatial distribution of selected genomic loci and their epigenetic states, yet remain limited by low detection efficiency and specialized high-content imaging instrumentations^17^. Thus, despite these important advances, a broadly accessible method that combines sub-micron-scale spatial precision, minimal custom hardware and software dependence, and compatibility with intact cells and tissues has yet to be achieved.

Here, we introduce ECHOS (Epigenetic CUT&Tag via High-resolution Optical Selection), a spatial chromatin profiling platform that integrates high-resolution imaging and high-throughput sequencing to enable precise, spatially targeted epigenetic characterization across biological scales. In ECHOS, Tn5 transposes deposit photo-caged adapters into DNA sequences associated with an epigenetic mark, rendering all tagmented DNA inert to amplification until the adapters are uncaged at user-defined regions of interest (ROIs) with a near ultraviolet (UV) laser light. We benchmarked the ability of ECHOS to capture the DNA-binding profile of the Lamin protein as well as multiple types of histone modifications in cells and tissues with ChIP-seq and CUT&Tag, showing comparable capture efficiency and specificity. We then further enhanced the sensitivity of ECHOS by adopting the strategy of linear amplification (ECHOS+), making it feasible to characterize the histone modification landscape of subcellular chromatin domains. Using ECHOS+, we revealed distinct gene regulatory logics at different layers of human ectocervical epithelium. We also showed that the chromatin in micronuclei (MN) exhibited a different H3K27ac profile compared to intact primary nuclei, suggesting an epigenetic dysregulation of the chromosome fragments in MN. Lastly, we found that female aging altered the H3K27me3 and H3K27ac landscape of the inactive X chromosome (Xi), especially at the regulatory regions of a subset of genes escaping X chromosome inactivation in human primary skin fibroblasts, shedding light on how aging contributes to sex-biased cellular phenotypes.

## Results

### The ECHOS Workflow

To enable precise spatial epigenome profiling of user-defined ROIs, we built upon our previous work in sequencing DNA of fixed samples at sub-micron (∼300 nm) resolution using an imaging-guided, laser-targeted photo-uncaging protocol^18^. The workflow of ECHOS consists of four major stages: (1) construction of a photocaged DNA library in situ; (2) selective uncaging of the DNA molecules using targeted illumination with a 405-nm laser light; (3) sample digestion and DNA purification; and (4) library preparation followed by sequencing (**Fig. 1A**).

**Figure 1.**
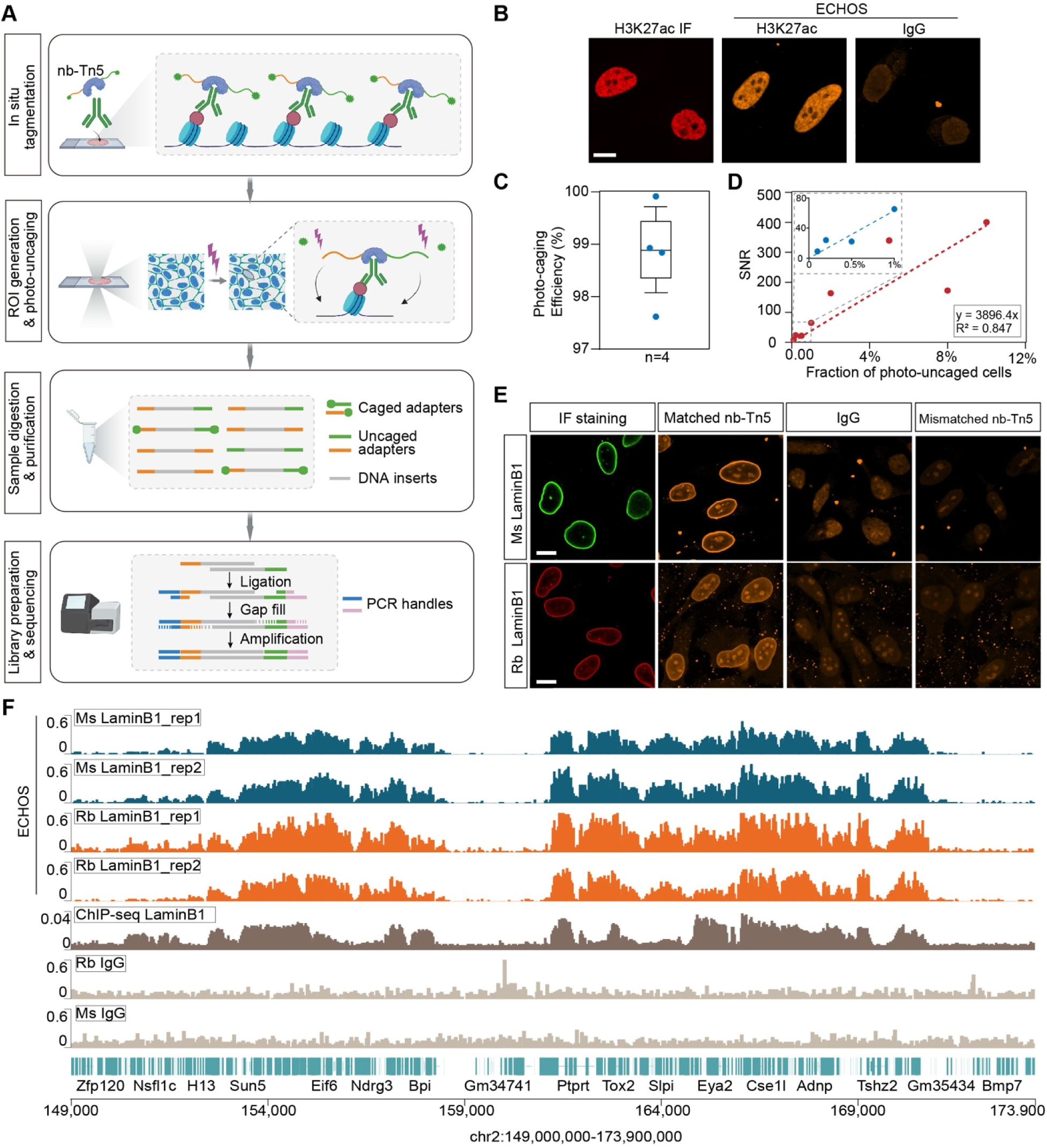
Overview and validation of the ECHOS workflow. **A** Schematic of the ECHOS workflow. Primary antibodies and nb–Tn5 are applied in situ, followed by ROI selection and photo-uncaging using targeted 405-nm laser illumination. Only photo-uncaged DNA fragments are competent for downstream ligation and amplification. **B** The H3K27ac ECHOS library showed the same nuclear pattern as the H3K27ac immunofluorescence (IF). Scale bars, 10 μm. **C** Photo-caging efficiency of ECHOS measured across independent biological replicates (n = 4). **D** Signal-to-noise ratio (SNR) as a function of the fraction of cells photo-uncaged. Each point represents an independently prepared ECHOS library. Dashed line indicates linear fit (slope = 3,896.4; R² = 0.847). Inset: expanded view of the low-fraction range. **E** Validation of the spatial specificity of ECHOS using Lamin B1 as a representative nuclear envelope target. Scale bars, 10 μm. **F** Genome browser tracks comparing ECHOS libraries generated using mouse (Ms) or rabbit (Rb) anti-Lamin B1 antibodies, benchmarked against a public ChIP–seq dataset.

First, fixed cells or tissue sections are stained with a primary antibody recognizing a target epigenetic mark such as a histone modification or a DNA-binding protein. A nanobody–Tn5 transposase fusion protein (nb-Tn5) from a species that matches that of the primary antibody is then introduced to the sample to enable the in situ insertion of photocaged DNA adaptors into chromatin regions at or near the epigenetic mark sites. The nb-Tn5 protein-mediated genome transposition have recently been demonstrated to enable genome-wide chromatin profiling^13,19^. The photo-caged adaptors contain fluorophores that are connected to the 5’ end of the DNA via photocleavable linkers (sequence information in **Table S1**), allowing the visualization of the tagmented chromatin. Second, ROIs are defined based on their anatomical or subcellular features with confocal imaging. Depending on the application, ROIs can be selected manually or identified through automated image segmentation workflows. A commercially available photo-stimulation module (the same hardware that is used in the fluorescence recovery after photobleaching [FRAP] assay) is then used to automatically apply focused 405-nm laser to selectively uncage the adapters within the selected ROIs. This procedure cleaves off the fluorophores and exposes 5′ phosphate groups on the photo-uncaged adaptors for downstream ligation reactions. Third, following sample digestion and DNA purification, PCR handles are ligated to photo-uncaged adapters while photo-caged DNA fragments remain ligation-incompetent, ensuring that only DNA within the ROIs can be amplified. Finally, sequencing libraries are generated after gap filling and PCR amplification and are sequenced on the Illumina platform.

### Validating the ECHOS Workflow

We first assessed the efficiency of ECHOS’ photo-caging mechanism by capturing the H3K27ac profiles of HeLa cells (**Fig.1B**). Briefly, following in situ tagmentation, DNA fragments were isolated from the cells and split into halves. One half was exposed to the 405 nm laser light to photo-uncage the inserted adaptors, while the other half was protected from light. The two halves were then subject to the rest of the ECHOS workflow, and the library sizes (also known as library complexity, defined as the number of unique sequence-able fragments after de-duplication) of the pair were compared. Across four independent experiments, ECHOS achieved on average approximately 98% photo-caging efficiency **(Fig.1C)**, demonstrating its high signal-to-noise ratio (SNR).

Next, we sought to further quantify the SNR for ECHOS experiments. Given the noise mechanism, we expected the amount of noise to scale with the total number of cells in the sample (and correspondingly the total amount of fragmented DNA), while the amount of signal is proportional to the area of the ROIs. Therefore, we calculated the relationship between the SNR and the fraction of cells photo-uncaged in situ, or equivalently, the fraction of the total ROI area that is photo-uncaged. To do this, we selected various proportions of the cultured HeLa cells and measured the resulting library sizes. As expected, the SNR increased almost linearly with the fraction of selected cells (R² = 0.85; **Fig.1 D**). We found that the SNR is approximately 5,396 times the fraction of cells selected. Notably, ECHOS produced high-quality libraries with SNR >9 even when only 0.1% of the sample (equivalent to ∼5 cells) was photo-uncaged.

Lastly, we examined the spatial specificity of ECHOS by targeting Lamin B1 protein-associated chromatin due to its stereotypic localization at the nuclear periphery. We observed nb-Tn5–mediated localization of fluorescent photo-caged adaptors at the nuclear periphery only when the anti-Lamin B1 primary antibody and nb-Tn5 were from the same species (**Fig. 1E**). Using the mouse nb-Tn5 against the rabbit anti-Lamin B1 antibody or using the rabbit nb-Tn5 against the mouse anti-Lamin B1 antibody abolished the specific localization (**Fig. 1E**). Using IgG instead of the primary antibody also prevented the localization of the photo-caged adaptors to the nuclear periphery (**Fig. 1E**). Sequencing of these Lamin B1-targeted ECHOS libraires yielded reproducible signal coverage profiles (**Fig. 1F**). These ECHOS profiles were also highly consistent with that of a publicly available Lamin B1 ChIP-seq data^20^ (**Fig. 1F & S1**), confirming the spatial accuracy of ECHOS.

### Benchmarking the Performance of ECHOS

To further assess the ability of ECHOS to accurately capture the epigenetic landscape of chromatin, we obtained the H3K4me3 profile of HeLa cells using ECHOS (**Fig. 2A**) and CUT&Tag, respectively. We also obtained publicly available H3K4me3 ChIP-seq datasets generated on HeLa cells from the ENCODE database^21^. Genome-wide correlation analysis revealed that the H3K4me3 ECHOS coverage profiles were highly reproducible across biological replicates (Pearson r >0.9) and were in strong concordance with both CUT&Tag and ChIP-seq profiles (**Fig. 2B**). Furthermore, more than 90% of ECHOS H3K4me3 peaks overlapped with ChIP-seq (**Fig. 2C**) and CUT&Tag peaks (**Fig. 2D**), respectively. Finally, fraction of reads in peaks (FRiP) analysis showed that ECHOS achieved signal enrichment at a level comparable to CUT&Tag and substantially higher than ChIP-seq **(Fig. 2E)**. It is worth mentioning that the ECHOS H3K4me3 libraries were generated using only ∼5,000 input cells, whereas both CUT&Tag and ChIP-seq require at least one order of magnitude more cells as starting materials.

**Figure 2.**
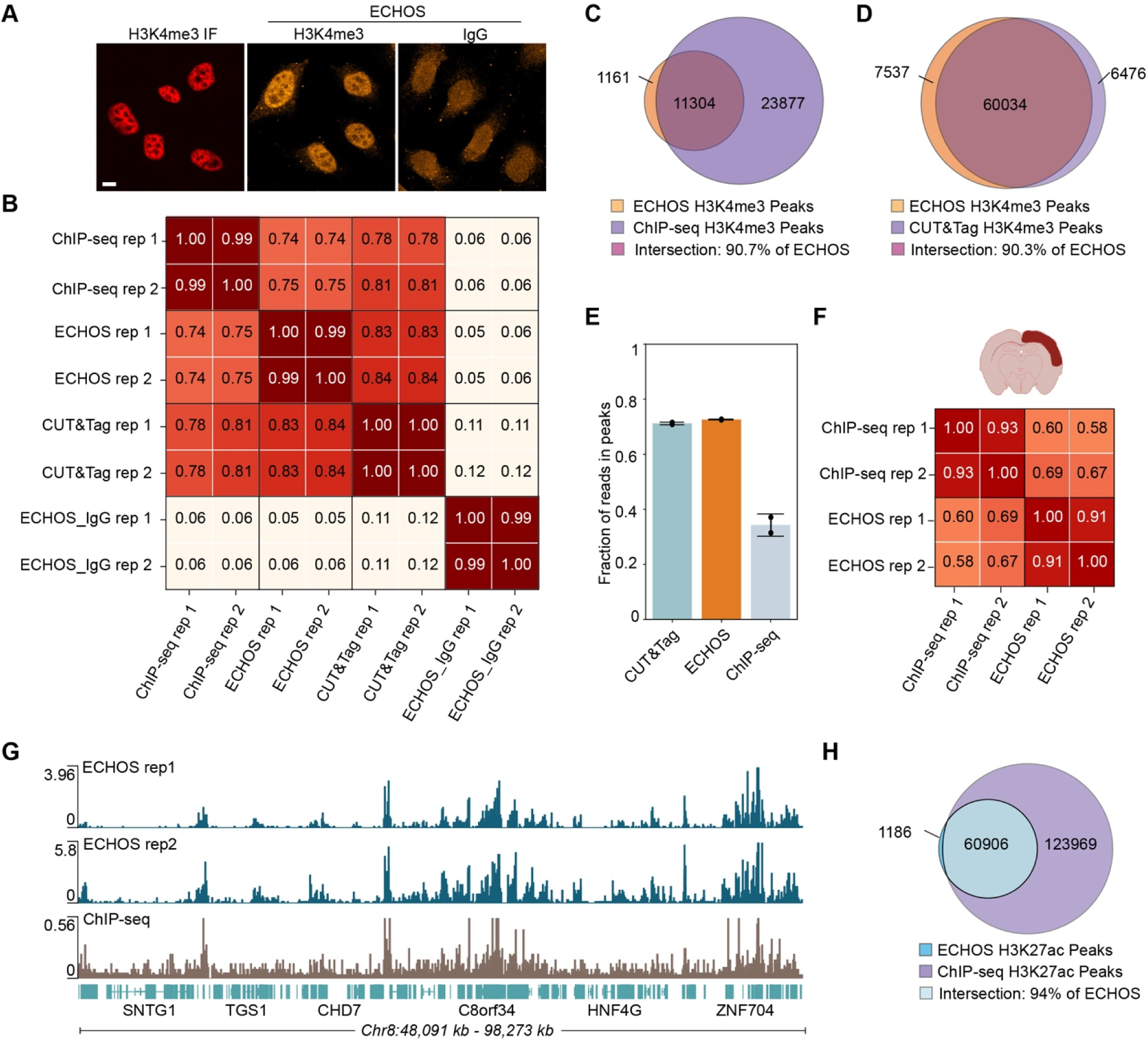
ECHOS recapitulates reference histone modification landscapes in cells and tissues. **A** Representative confocal images showing IF staining and ECHOS libraries for H3K4me3 in cultured HeLa cells. Scale bars, 10 μm. **B** Pairwise Pearson correlation matrix computed from genome-wide 10-kbp binned coverage tracks for H3K4me3 datasets generated by ChIP–seq, CUT&Tag, and ECHOS, respectively. **C** Overlap between ECHOS and ChIP–seq H3K4me3 peaks in HeLa cells. Venn diagram shows the number of peaks uniquely identified by each method and the shared peak set. **D** Overlap between ECHOS and CUT&Tag H3K4me3 peaks. Venn diagram shows the number of peaks unique to each method and the shared set. **E** Fraction of reads falling within called peaks for H3K4me3 libraries generated by CUT&Tag, ECHOS, and ChIP–seq, respectively. Two biological replicates for each method were used. **F** Pairwise Pearson correlation matrix computed from genome-wide 10-kbp binned coverage tracks for H3K27ac profiles generated from mouse cerebral cortex by ChIP–seq and ECHOS, respectively. **G,** Genome browser tracks showing representative H3K27ac coverage profiles in mouse cerebral cortex for ECHOS and ChIP–seq data, respectively. **H,** Overlap between ECHOS and ChIP–seq H3K27ac peaks in mouse cerebral cortex. Venn diagram shows 94% of ECHOS H3K27ac peaks intersect with ChIP–seq peaks.

Next, we tested the performance of ECHOS in the context of tissues. We applied ECHOS to brain sections from adult mice and captured the H3K27ac profiles of nuclei in the left cerebral cortex. Genome-wide coverage correlation analysis of ECHOS data showed strong reproducibility between biological replicates (Pearson r = 0.91; **Fig. 2F**). The coverage profiles of ECHOS were similar to that of a public ChIP-seq dataset from mouse cerebral cortex (**Fig. 2G**). Indeed, 94% of ECHOS H3K27ac peaks were also found in the ChIP-seq dataset (**Fig. 2H**).

Together, these results show that ECHOS captures epigenetic profiles of cells and tissues at a level of accuracy that matches the accuracy of the standard methods.

### Sensitivity Enhancement with ECHOS+

Encouraged by the performance of ECHOS, we went on to further improve its sensitivity to overcome the challenge of ultra-low input when it comes to epigenetic profiling of small subnuclear regions.

In ECHOS, the nb-Tn5 inserts the photo-caged forward and reverse adapters in random orientations and roughly half of the tagmented genomic DNA do not have adapters in the correct orientation to be later amplified by PCR. Furthermore, PCR-based library preparation is extremely sensitive to size variations of the DNA fragments, with shorter DNA fragments prone to be overamplified by PCR and over-clustered during Illumina sequencing compared to longer fragments. In contrast, linear amplification has been used to circumvent amplification and sequence composition bias caused by PCR and is particularly beneficial when only limited amount of input DNA is available. In fact, multiple genome and epigenome sequencing methods have transitioned to linear amplification for increased sensitivity^22–25^. Therefore, we hypothesized that adapting ECHOS to use linear amplification would greatly improve its sensitivity for subcellular level epigenetic profiling.

To test this hypothesis, we designed ECHOS+ by merging ECHOS with an existing Tn5 transposition-based linear amplification protocol^24,25^. In ECHOS+, nb-Tn5 inserts only one type of photo-caged adaptors into the genomic DNA at or near the sites of an epigenetic mark. This adaptor contains a non-functional, partial T7 promoter sequence at the 5’ end and is also connected to a fluorophore via the photocleavable linker (sequence information in **Table S1**). Upon photo-uncaging, the fluorophores on the adaptors are cleaved off to expose 5’ phosphate groups, allowing the subsequent ligation of the remaining sequences of the T7 promoter. Thus, only DNA fragments subject to photo-uncaging within the ROIs will contain functional, full-length T7 promoter sequences to permit downstream linear amplification (**Fig. 3A**). In vitro transcription (IVT) by T7 RNA polymerase creates ∼1,000-fold RNA copies of insertion sites. Following reverse transcription (RT), second-strand synthesis, and cDNA fragmentation, the resulting amplicons are prepared for sequencing by attaching index sequences. Because ECHOS+ relies on only the partial T7 promoter–containing adapters (rather than forward and reverse adapters in ECHOS), the issue of efficiency loss due to mismatched adapter orientations and varying distances between insertions should be mitigated. Additionally, linear amplification by IVT will ensure high fidelity and uniformity since short fragments are not over-represented in the sequencing libraries.

**Figure 3.**
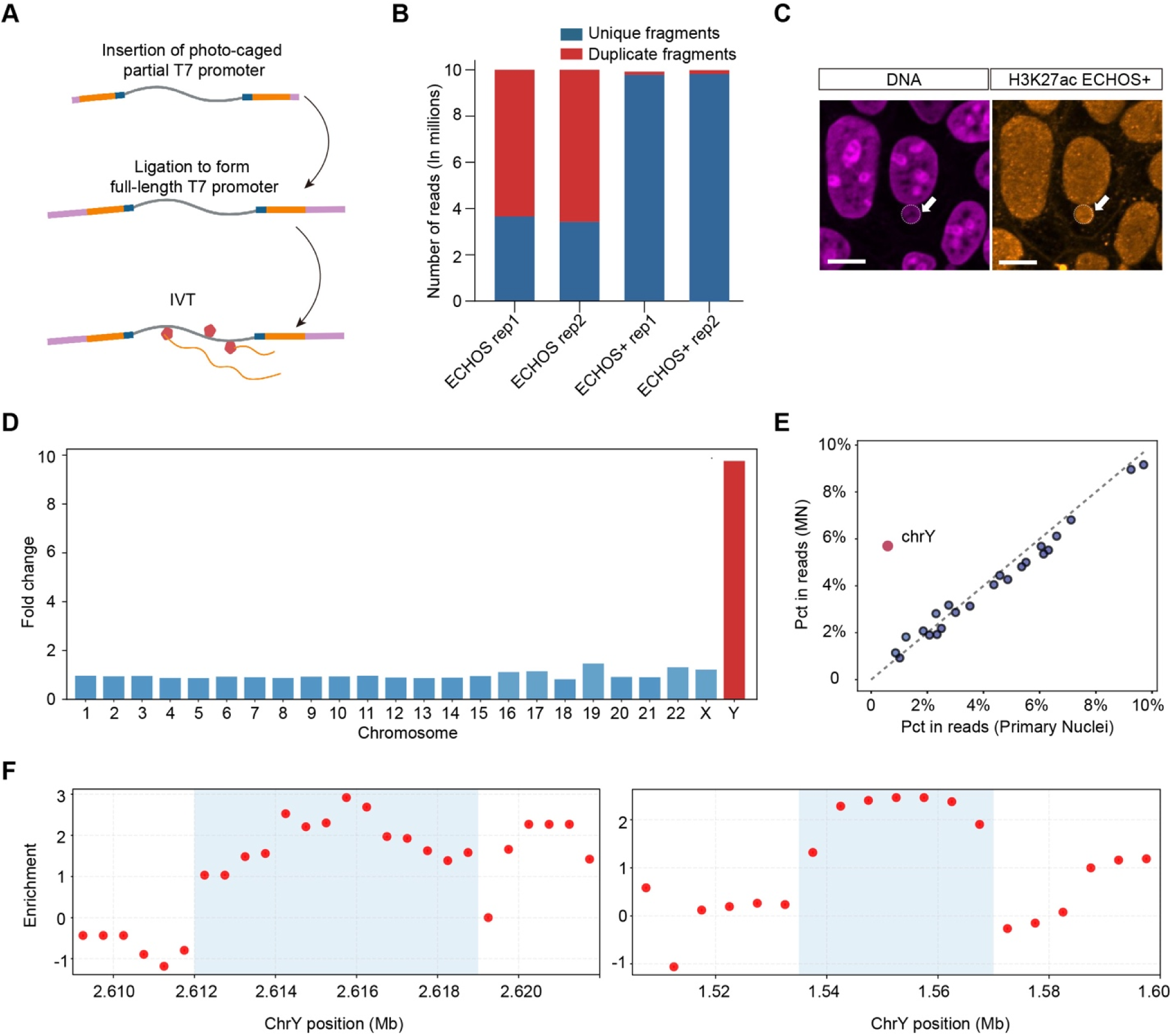
ECHOS+ captures the epigenetic profiles of micronuclei (MN). **A** The linear amplification step in ECHOS+ workflow. **B** Comparison of library complexity between ECHOS and ECHOS+ H3K27ac datasets. Stacked bars show the number of unique and duplicate fragments. ***C*** Confocal imaging of MN (left) and H3K27ac ECHOS+ library (right). White arrow denotes MN. Scale bars, 10 μm. **D** Fold change of mapped sequencing reads for each chromosome in MN ECHOS+ libraries over primary nucleus ECHOS+ libraries. **E** Percentage of each chromosome in sequencing read space in MN ECHOS+ libraries and primary nucleus ECHOS+ libraries. **F** Representative Y chromosome regions with elevated H3K27ac signals in MN over primary nuclei.

Indeed, despite the design changes, ECHOS+ maintains a high photo-caging efficiency (**Fig. S2A**) and SNR (**Fig. S2B**). By comparing the H3K27ac libraries of HeLa cells generated by ECHOS and ECHOS+, we found that ECHOS+ generated approximately 2.5 times more unique fragments than ECHOS at the same number of input cells and sequencing depth (**Fig. 3B**). Besides recovering most of the H3K27ac peaks identified by ECHOS, ECHOS+ captured an additional set of peaks that had no or limited coverage in ECHOS data (**Fig. S2C&D**).

Together, these results suggest that ECHOS+ significantly improves the dynamic range of signal detections.

### ECHOS+ Enables Epigenetic Profiling of Micronuclei (MN)

We next evaluated the ability of ECHOS+ to capture the epigenetic profiles of subcellular compartments by focusing on MN. MN contain mis-segregated chromosomes and are spatially separated from the primary nucleus^26^. They are often found in cancer cells and can propel tumor evolution through genetic, epigenetic, and transcriptional changes^27,28^. Here, we leveraged an engineered cellular system in which MN can be generated by selectively inducing the mis-segregation of human Y chromosome by drug treatment^29^. Thus, this system generates Y chromosome-containing MN (hereafter referred to as Y-MN) and provides ground truth sequence information for the validation of ECHOS+.

We applied ECHOS+ to characterize the H3K27ac landscape of Y-MN (**Fig. 3C**) as well as primary nuclei of untreated cells (hereafter referred as control). Encouragingly, we found a significant enrichment of sequencing reads that were mapped to the Y chromosome in the Y-MN ECHOS+ libraries over the control libraries (**Fig. 3D**). This level of Y chromosome read enrichment was consistent with the result of a previous study in which Y-MN were isolated and sequenced^30^. Furthermore, the levels of read coverage for the rest of the non-Y chromosomes remained unchanged between the Y-MN and the control data (**Fig. 3E**), suggesting that ECHOS+ selectively targeted Y-MN with high accuracy.

We next compared the H3K27ac landscape between Y chromosomes in the Y-MN and the same Y chromosome regions in the control. Analysis using normalized coverage data indicated differential H3K27ac signals between Y-MN and the control as we identified multiple Y chromosome domains whose H3K27ac levels were elevated in Y-MN vs. the control (Methods) (**Fig. 3F**), suggesting that MN chromatin is subject to different epigenetic regulations from the chromatin in primary nuclei.

Together, our data establish that ECHOS+ can resolve the epigenetic landscapes of subcellular level DNA compartments like the MN.

### ECHOS+ Enables Compartment-Specific Epigenetic Profiling in Human Ectocervical Epithelium

After validating ECHOS+, we went on to demonstrate its usage in facilitating the understanding of gene regulatory mechanisms under various biological context.

We first turned to the human cervical epithelium especially the ectocervical epithelium because it serves as a barrier against microbial entry within the female reproductive tract^31^. A detailed molecular understanding of the ectocervical epithelial cell behavior and functions would have a profound impact on our effort in improving women’s reproductive health. The human ectocervical epithelium is a stratified squamous epithelium consisting of multiple layers: the basal and parabasal layers, followed by the intermediate and superficial layers^32^. For simplicity, we used ECHOS+ to capture the H3K4me3 profiles of two broad epithelial compartments using human cervical tissue slices: (1) a basal compartment, encompassing the basal layer and adjacent parabasal cells; and (2) a luminal compartment, composed of the intermediate and superficial layers (**Fig. 4A**). Focusing on the H3K4me3 signals at the promoter regions, we identified genes which showed differential promoter H3K4me3 signals between the two epithelial compartments (**Fig. 4B**). Some of the genes which have been previously shown to be associated with the functions of the cervical epithelium, such as *MUC21*^33,34^ and *GRHL1*^35,36^, were shown to have elevated promoter H3K4me3 signals at the luminal compartment (**Fig. 4B**).

**Figure 4.**
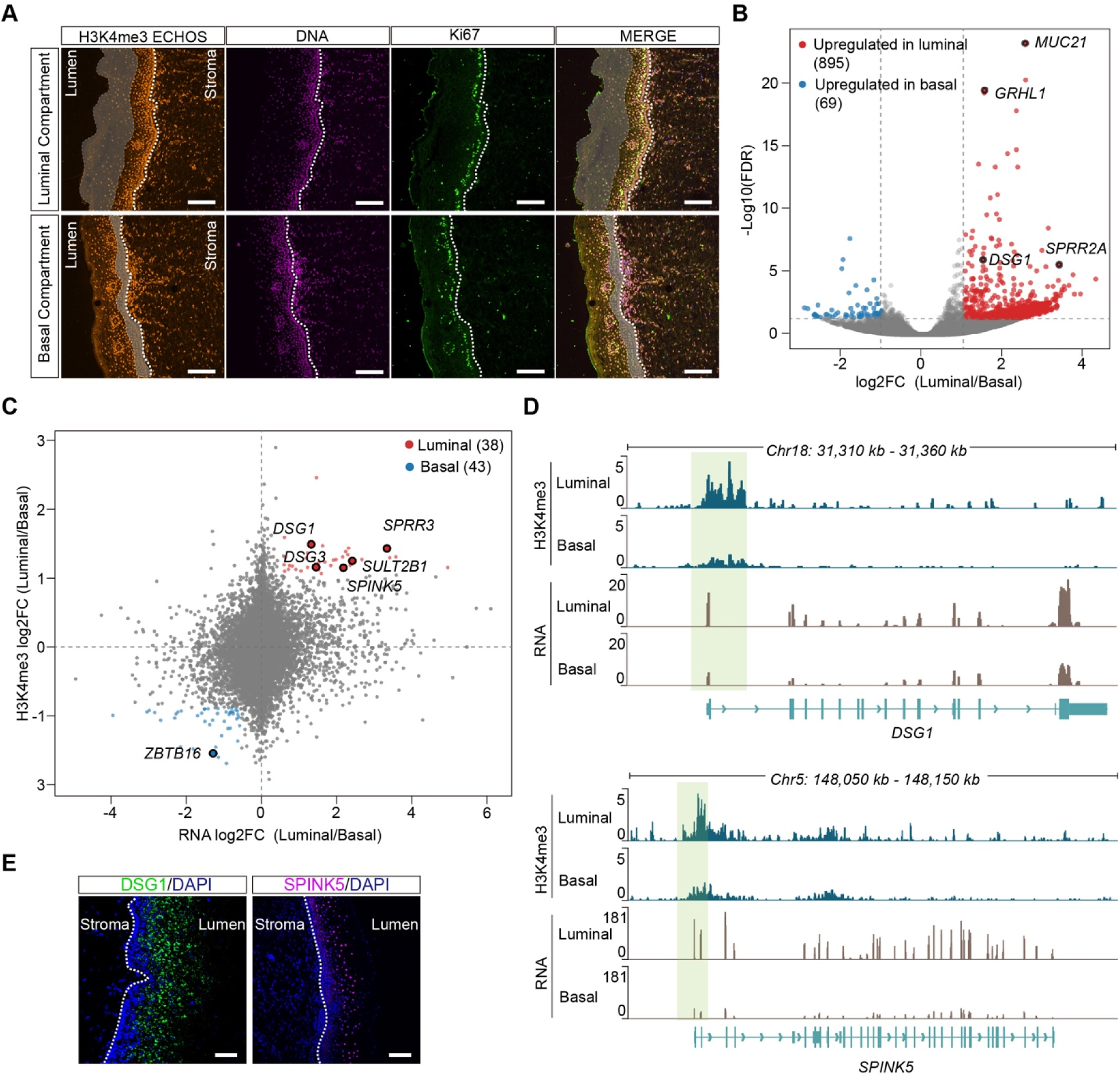
ECHOS+ resolves compartment-specific epigenetic states in human ectocervical epithelium. **A** Representative confocal imaging of human ectocervix tissue sections showing the basal and luminal compartments for photo-uncaging (shaded areas). Ki67 staining highlights the proliferative cell layer of the epithelium. The white dashed lines mark the boundary between the epithelium and the stroma. Scale bars, 100 μm. **B** Genes with differential promoter H3K4me3 signals between luminal and basal compartments profiled by ECHOS+. **C** Joint analysis of compartment-specific H3K4me3 promoter signals and matched gene expression levels. **D** Genome browser tracks showing H3K4me3 profiles and matched RNA expression for DSG1, and SPINK5. Green shaded regions indicate promoter-proximal H3K4me3 signals. **E** RNA FISH validation of luminal compartment-enriched expression patterns of DSG1 and SPINK5. The white dashed lines mark the boundary between the epithelium and the stroma. Scale bars, 100 μm.

To examine how changes in the H3K4me3 levels between the two epithelial compartments impact gene expression, we performed spatial transcriptomics analyses to capture the RNA expression profiles of the two compartments using a recently developed technology called PHOTON^37^. We then plotted the fold change (luminal vs. basal) of the promoter H3K4me3 level of each gene against its RNA expression fold change (**Fig. 4C**), which identified a set of genes with positive correlations. For example, the promoter regions at *DSG1* and *SPINK5* displayed stronger H3K4me3 signals in the luminal compartment. Correspondingly, the RNA expression levels of *DSG1* and *SPINK5* were also significantly higher in the luminal compartment (**Fig. 4D**). The luminal compartment-enriched spatial expression patterns of *DSG1* and *SPINK5* were further confirmed by RNA fluorescent in situ hybridization (RNA FISH) (**Fig. 4E**). Of interest, *DSG1* has recently been implicated in the human ectocervical epithelium functions^38^, suggesting that ECHOS+ can be leveraged to identify functional relevant genes.

Thus, these data demonstrate the effectiveness of ECHOS+ to extract spatially resolved epigenetic profiles from clinical samples like the human cevix.

### ECHOS+ Captures Aging-induced Epigenetic Changes on Inactive X Chromosomes (Xi)

Lastly, we leveraged ECHOS+ to explore how human aging impacts the epigenetic landscape of chromatin. To this end, we stained H3K27me3 in human primary skin fibroblasts collected from both young (10-25 years old) and aging (>70 years old) healthy female donors. Intriguingly, we noticed a nuclear focus with a strong H3K27me3 fluorescence intensity in both the young and aging cells (**Fig. 5A**). To reveal the sequence identity of this focus, we performed ECHOS+ targeting H3K27me3 within the focus. Correspondingly, we also performed ECHOS+ targeting H3K27me3 of the whole nucleus of the same fibroblast line as a control. Interestingly, we observed a strong enrichment of sequencing reads that were mapped to the X chromosome in the H3K27me3-positive nuclear focus libraries, but not in the control libraries (**Fig. 5B&S3A**). This suggests that the H3K27me3-positive nuclear focus contained mostly the X chromosome. Given that H3K27me3 is a repressive histone mark, we hypothesized that the nuclear focus contained the Xi. RNA FISH of the *XIST* long non-coding RNA showed a strong overlap between the *Xist* signals and the nuclear focus (**Fig. 5C**), confirming our hypothesis. In fact, previous work has shown that during female embryonic development, the Xi condenses into a compact structure called the Barr body^37^. Thus, ECHOS+ accurately captured the sequence information of the Barr body.

**Figure 5.**
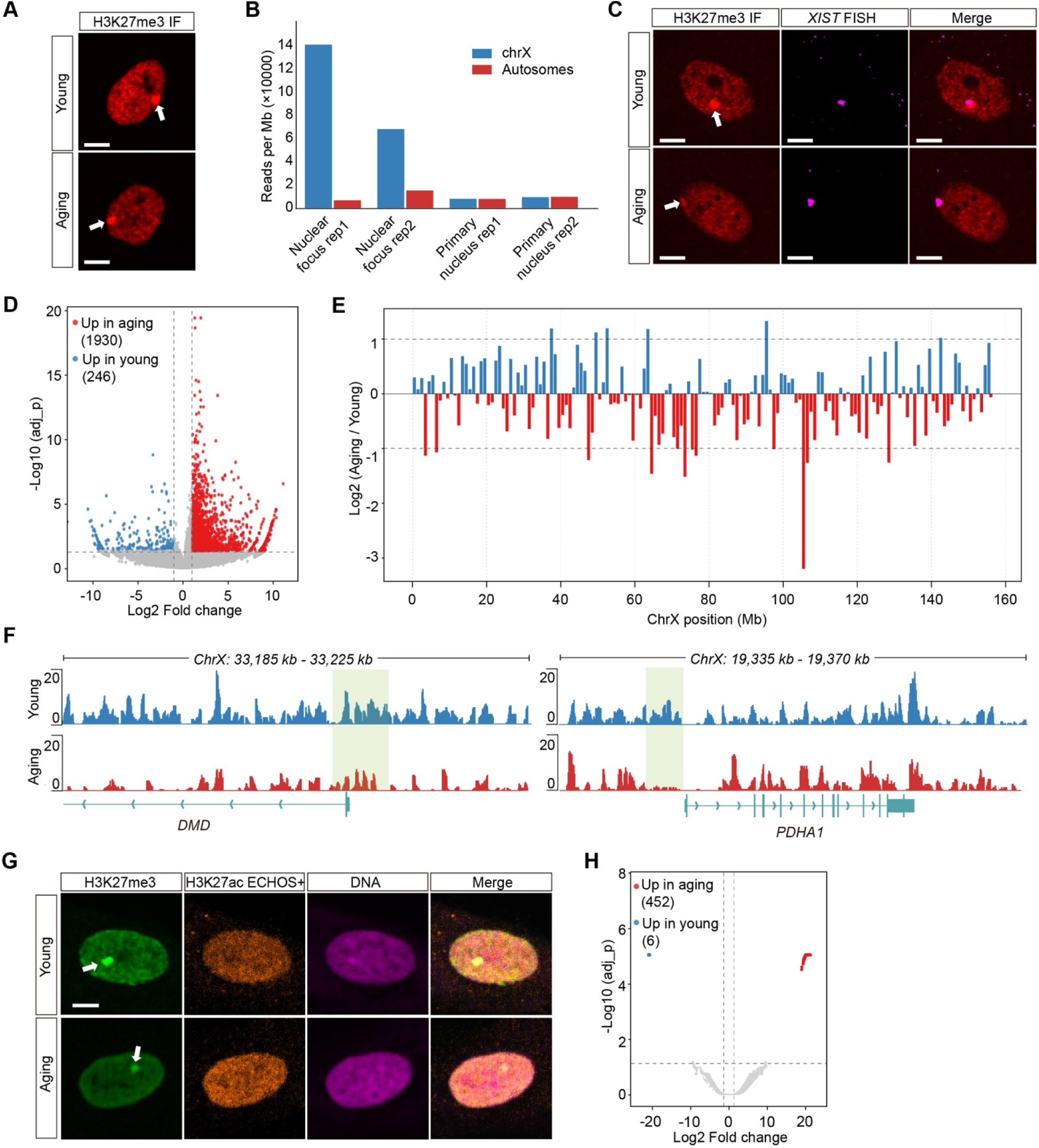
ECHOS+ reveals age-associated epigenetic alterations on Xi **A** Representative H3K27me3 IF images of young and aging human female skin fibroblasts. The white arrow points to a prominent H3K27me3-dense focus in the nucleus. Scale bars, 10 µm. **B** Quantification of normalized H3K27me3 ECHOS+ sequencing reads mapped to the X chromosome versus autosomes across indicated groups. **C** Representative images showing the colocalization between the H3K27me3-positive nuclear focus (white arrow) and XIST RNA. Scale bars, 10 µm. **D** Volcano plot showing differential H3K27me3 peaks on young and aging Xi. **E** Normalized Xi H3K27me3 signals in aging cells over young cells. **F** H3K27me3 genome browser views of representative loci (DMD and PDHA1) in both the young and aging group. **G** Representative images of H3K27me3 IF stain together with in situ H3K27ac ECHOS+ library of young and aging human female skin fibroblasts. The white arrow denotes the Barr body. Scale bars, 10 µm. **H** Volcano plot showing differential H3K27ac peaks on young and aging Xi.

We then asked if aging alters the H3K27me3 landscape of the Xi. We first measured the H3K27me3 fluorescence intensity at the Barr body of the young and aging fibroblasts and did not see significant differences (**Fig. S3B**), which may be due to cellular and patient level heterogeneity. Next, we compared the Barr body-specific H3K27me3 ECHOS+ data between the two age groups. Of interest, we identified differential H3K27me3 peaks (**Fig. 5D**), showing that aging could induce both site-specific gains and losses of H3K27me3 signals on Xi. Visualization of the fold change in H3K27me3 signals in the aging group over the young group along the coordinates of the Xi showed pronounced region-to-region signal variations (**Fig. 5E**). This highlights the ability of ECHOS+ to resolve nuanced epigenetic changes that are undetectable by fluorescence microscopy alone. Notably, certain loci of Xi that showed weakened H3K27me3 signals in the aging group overlapped with or neighbored several known X inactivation escape-prone or structurally labile genes such as *DMD*, *PDHA1*, and *ARHGAP6*^37,39^. Detailed examination of these three gene loci showed reduced H3K27me3 signals at the promoter regions (**Fig. 5F&S3C**).

To further understand how aging alters the epigenetic landscape of the Xi, we obtained the H3K27ac profiles of the Xi in the young and aging fibroblasts using ECHOS+ (**Fig. 5G**). Differential peak analysis identified significantly more upregulated H3K27ac peaks in the aging fibroblasts than in the young ones (**Fig. 5H**). Notably, none of the 452 upregulated H3K27ac peaks on the Xi of the aging fibroblasts were located within the 5-kbp window of a transcription start site. Rather, these peaks were found in distal regulatory regions, suggesting a different layer of epigenetic regulation from H3K27me3 during female aging.

Finally, qPCR analysis of the expression level of *XIST* showed no difference between the two age groups (**Fig. S3D**), suggesting that mechanisms other than *XIST* dysregulation may mediate these aging-induced epigenetic changes at Xi.

Together, these data demonstrate the ability of ECHOS+ to accurately capture the epigenetic alterations of a subnuclear structure during pathological conditions such as aging.

## Discussion

Despite recent advances in spatial omics technology development, spatially resolved profiling of the epigenome remains technically challenging, particularly when the measurements need to extend to micron-scale nuclear structures. Here, we address this challenge by developing ECHOS/ECHOS+, a platform that seamlessly integrates high-resolution imaging with high-throughput sequencing. This platform fills the critical gap in the spatial omics toolkit development when it comes to sensitive epigenetic profiling across length scale—from tissue regions to subnuclear compartments.

At the tissue level, we showed that ECHOS recovered a significant amount of H3K27ac signals of the adult mouse cerebral cortex identified by ChIP-seq (**Fig. 2F-H**). However, unlike ChIP-seq which would require a series of steps to isolate thousands of cells from the brain tissue, ECHOS achieved a similar level of signal coverage by photo-uncaging the cerebral cortex region of a thin mouse brain slice in situ. This is especially advantageous for clinical research in which obtaining a large quantity of input cells for ChIP-seq is often not feasible. Indeed, leveraging ECHOS+, we obtained region-specific H3K4me3 profiles in the human ectocervical epithelium using only a few thin tissue slices (**Fig. 4**), demonstrating the potential compatibility of our platform with routine clinical sample acquisition and preparation.

At the subcellular level, we first applied ECHOS+ to target MN. MN represent a formidable challenge to conventional epigenetic profiling methods due to their minimal DNA content and small sizes. Indeed, previous attempts to purify MN required many input cells, might disrupt fragile MN membranes, alter their chromatin composition, or introduce contamination^40^. In contrast, ECHOS+ enabled direct in situ epigenetic profiling of MN within intact cells and generated high-complexity libraries from only a few thousand cells. Recent work using immunofluorescence (IF) and mass spectrometry has uncovered reductions in several histone modifications such as H3K9ac and H3K27ac in MN when compared with primary nuclei in several cell lines^28^, suggesting epigenetic mis-regulation of miss-segregated chromosomes. Although we did not observe a global reduction of H3K27ac level in Y-MN likely due to differences in cellular context (**Fig. 3C**), we uncovered heterogeneous, domain-scale redistributions of H3K27ac along the Y-MN chromatin (**Fig. 3F**). These changes in MN histone modifications may act as a biochemical signal that affects downstream processes such as the sensing of cytosolic double-stranded DNA by the cGAS–STING pathway.

Furthermore, by taking advantage of the spatial structure of the Barr body and the high spatial resolution of ECHOS+, we successful captured the epigenetic profiles of Xi directly from intact cells. We found that the both the H3K27me3 and H3K27ac landscapes at Xi were altered in aging human skin fibroblasts (**Fig. 5D-H**), and some of the Xi genome regions with dampened H3K27me3 signals coincided with known X inactivation escape-prone or structurally labile genes such as *DMD*, *PDHA1*, and *ARHGAP6* (**Fig. 5F&S2D**). These observations suggest a potential aging-dependent gene regulatory mechanism acting on the Xi. Indeed, recent studies of the aging female mice showed that aging preferentially changed gene expression on the X chromosome relative to autosomes, with select genes on the Xi undergoing activation^41,42^. Furthermore, increased chromatin accessibility was discovered at the Xi, but not at the autosomes or the active X chromosome in aging female mouse kidney^42^. Together, these data start to paint a picture of how epigenetic alterations of the Xi may be linked to sex-specific aging phenotypes.

Looking ahead, going beyond stable nuclear structures, ECHOS+ is poised to address important questions related to the membrane-less, phase-separated phenomenon known as biomolecular condensates, such as the nucleoli, nuclear speckles, and transcriptional condensates^43^. These structures are difficult to isolate biochemically without disrupting their native interactions. In contrast, ECHOS+ could uniquely dissect the epigenetic logic governing the formation and function of these nuclear condensates in situ.

Although ECHOS and ECHOS+ represent substantial advances in high resolution spatial epigenome profiling, several limitations exist. First, during photo-uncaging with the laser light, the laser passes through the entire thickness of the cell and can potentially uncage DNA fragments above and below the focal plane outside of the targeted ROIs. Currently, we navigate this effect by optimizing the optical conditions during imaging to make sure that most of the ROIs in a field of view (FOV) stay within the focal plane. Nevertheless, both the axial and lateral resolution could be further improved by targeting the ROIs with two-photon absorption, enabling fully volumetric photo-uncaging. Second, the current setup of the photo-stimulation module requires the photo-uncaging process be performed in a pixel-by-pixel, ROI-by-ROI manner, limiting the experimental throughput. Future iterations incorporating new photo-stimulation hardware such as a digital mirror device would allow photo-uncaging of all ROIs within a FOV simultaneously. Third, ECHOS and ECHOS+ currently work best with fresh frozen tissue samples. Extending their use to formalin-fixed paraffin-embedded samples would unlock epigenomic analysis across retrospective patient cohorts, providing powerful opportunities to study long-term disease progression, therapeutic response, and patient-specific gene regulatory alterations. Finally, ECHOS and ECHOS+ only capture one epigenetic mark at a time. Enabling simultaneous profiling of multiple epigenetic marks as well as other modalities such as the transcriptome would provide a more comprehensive view of the gene regulatory landscape.

In summary, ECHOS/ECHOS+ serve as a powerful platform for resolving epigenetic landscapes across different biological scale. By combining imaging-and sequencing-based readouts, this platform will uncover new connections at the interface of physical and epigenomic space.

## Methods

### Tissue collection

Whole brains were collected from 8–12 weeks old adult female C57BL/6J mice (The Jackson Laboratory). Immediately after dissection, brains were briefly rinsed in cold PBS, blotted to remove excess moisture, and embedded in OCT compound (Tissue-Tek, 4583 Sakura). Tissue blocks were samp frozen in liquid nitrogen, kept on dry ice, and stored at –80°C until sectioning. All procedures were performed with prior approval of the UT Southwestern Medical Center on Use and Care of Animals, in accordance with the guidelines established by the National Research Council Guide for the Care and Use of Laboratory Animals. Mice were housed in the UT Southwestern Medical Center animal facility, in an environment controlled for light (12 h on/off) and temperature (21–23 °C) with ad libitum access to water and food.

A cervix tissue collection pipeline was established with an approved IRB through a collaboration with the Human Tissue, Biological Fluids, and Cell Culture Laboratory Core at UT Southwestern Medical Center. From gynecologic hysterectomy, full thickness ectocervical biopsies (1cm X 1cm) from premenopausal nonpregnant individuals were obtained. Exclusion criteria for individuals are women <21 years of age, the presence of active genitourinary infection at time of surgery, pathologic evidence of cervical dysplasia, or cervical cancer. Tissue samples were snap frozen in liquid nitrogen and embedded in OCT for ECHOS+ studies. Informed consent was obtained for study participants.

### Cell culture

HeLa cells (ATCC CRM-CCL-2) were cultured in DMEM, high glucose, pyruvate (Thermo Fisher Scientific, 11995073) supplemented with 10% heat-inactivated FBS (Thermo Fisher Scientific, 16140-071) and 1% Penicillin–Streptomycin (Gibco, 15140-122) and maintained at 37°C in 5% CO_₂_. Cells were routinely passaged using 0.05% Trypsin-EDTA. Primary human female skin fibroblasts were obtained from the Coriell Institute for Medical Research and were maintained under the same conditions as HeLa cells (DMEM, 10% heat-inactivated FBS, 1% Pen-Strep; 37°C, 5% CO_₂_). Cells were used between passages P6 to P10. For MN ECHOS+ experiments, a genetically engineered cell line (DLD-1 CENP-A replacement cell line) was obtained from the laboratory of Peter Ly (UT Southwestern Medical Center). Cells were maintained in DMEM supplemented with 10% FBS and 1% Penicillin–Streptomycin, with 200 μg/mL Geneticin, G418 Sulfate (Gibco, 10131035) included to maintain the integrated neomycin-resistance cassette. Induction of Y-MN was performed following the following a previously established scheme^29^. On Day 1, cells were seeded onto dishes; On Day 2, fresh growth medium containing 1 µg/mL doxycycline (DOX, Sigma-Aldrich, D3447) and 500 µM indole-3-acetic acid (IAA, Sigma-Aldrich, I5148) were added to induce degradation of endogenous CENP-A and expression of CENP-A^C-H3; On Day 3, DOX/IAA medium was replaced with fresh medium containing 100 ng/mL colcemid for 4 h to enrich mitotic cells with newly formed MN.

### Antibodies

The following primary antibodies were used in IF, CUT&Tag, ECHOS or ECHOS+ experiments: rabbit anti-H3K27me3 (Cell Signaling Technology, 9733), rabbit anti-H3K27ac (abcam, ab177178), rabbit anti-H3K4me3 (Cell Signaling Technology, 9751), mouse anti-SUZ12 (Santa Cruz Biotechnology, sc-271325), rat anti-Ki67 (Invitrogen, 14-5698-82), mouse anti-Lamin B1 (Proteintech, 66095-1-Ig), rabbit anti-Lamin B1 (Abcam, ab16048), mouse IgG1 isotype control (Cell Signaling Technology, 5415), and rabbit IgG isotype control (Proteintech, 30000-0-AP).

### The ECHOS workflow

Photo-caged adapters and the blocked mosaic end (ME) were obtained from Integrated DNA Technologies (IDT) and resuspended in ultrapure water (Invitrogen 10977015) to a final concentration of 100_µM. The ME was annealed to the adapters using the following reaction: 25_µM ECHOS_17, 25_µM ECHOS_21, 50_µM ECHOS_37, 10_mM Tris–HCl pH 8.0, 1 mM EDTA and 50_mM NaCl. The annealing reaction was heated to 95°C for 2 min in a thermocycler, and the temperature was gradually decreased from 20_°C over approximately 1_h. Annealed adaptors were mixed 1:1 with glycerol and stored at −20_°C until nb-Tn5 loading. A complete list of oligonucleotides sequences is provided in **Table S1**.

The plasmids encoding nb-Tn5 were gifts from New York Genome Center & Ivan Raimondi^13^. nb-Tn5 protein were produced in house and were loaded with annealed photo-caged adapter duplexes in 2x Tn5 loading buffer (20 mM Tris-HCl pH 7.5, 800 mM NaCl, 0.2 mM EDTA, 1 mM DTT, 60% glycerol) at a final adapter to Tn5 molar ratio of 8:1, and incubated at room temperature for 30–60 min.

To perform ECHOS on cultured cells, approximately 5000 cells were seeded onto Matrigel-coated 96-well glass-bottom plates (Cellvis P96-1.5H-N; Matrigel diluted 1:50 in DMEM). Following culture, cells were washed once with PBS (Gibco™ PBS, pH 7.4, 10010049) and fixed in 1% PFA freshly diluted from 16% Paraformaldehyde aqueous solution, EM grade (Electron Microscopy Sciences 15710) for 10 min at room temperature. Fixative was removed and samples were washed three times with PBS. Cells were permeabilized in 1% Triton X-100 (Sigma Aldrich, T9284-100ML) in PBS for 15 min at room temperature, followed by three PBS washes. To enhance the accessibility of nb-Tn5 to the chromatin, cells were treated in 0.1 N hydrochloride (HCl) for 5 min, then washed thoroughly with PBS. The same HCl treatment was also used in Epigenomic MERFISH^3^.

Samples were then blocked in 4% ultrapure BSA (ThermoFisher, AM2616) in PBS supplemented with 1% Triton X-100 for 1 h at room temperature. Following PBS washes, samples were incubated with primary antibodies overnight at 4 °C. After three PBS washes, samples were incubated with loaded nb–Tn5. For cell-based ECHOS experiments, loaded nb–Tn5 stock was diluted at 1:25 in freshly prepared Tn5 binding buffer (20 mM HEPES pH 7.5, 0.3–0.5 M NaCl, 0.2 mM EDTA, 0.01% digitonin (Sigma Aldrich, 300410), and 2 mM spermidine). Samples were incubated in the diluted, loaded nb–Tn5 for 1 h at room temperature and were protected from light. Unbound nb–Tn5 was removed by four washes in Tn5 binding buffer.

To initiate tagmentation, transposition buffer was prepared by supplementing Tn5 binding buffer with 10 mM MgCl_₂._ Cells were incubated in the transposition buffer for 1 h at 37 °C and were protected from light. After tagmentation, the reaction was stopped by incubating the cells with 20 mM EDTA and 10 mM Tris-HCl pH 8.0 for 10 min at room temperature, followed by PBS washes. Samples were then incubated with the nuclear stain DRAQ5™ (Abcam, ab108410, 1:1000 dilution in PBS) for 10 min at room temperature, washed twice with PBS, and immediately taken to the microscope for imaging.

Photo-uncaging of ROIs was performed on a Nikon A1R confocal microscope equipped with a 405-nm Galvo-XY photo-stimulation module (the same module used in the FRAP assay). For each relevant FOV within the sample, a multichannel confocal image was acquired to visualize the ECHOS library, sample morphology, and other protein markers. ROIs were defined either manually or algorithmically. Selective photo-uncaging was then performed by illuminating the ROIs with 405-nm laser light.

Following photo-uncaging, cells were digested with Proteinase K (NEB, P8107S) diluted (1:100) in reverse-crosslinking buffer (50 mM Tris-HCl pH 8.0, 50 mM NaCl, 0.2% SDS) for 1 h at 55 °C. Following digestion, DNA was purified using the NucleoSpin Gel and PCR Clean-Up kit (Takara, 740609) and eluted in 10–16 µL nuclease-free water. Ligation was then performed using pre-assembled indexed oligos (e.g., PSS_DNA_IDX_S501–S506) with T4 DNA ligase (NEB, M0202) in Quick Ligation buffer at 22 °C for 1 h. After ligation, PCR was performed using Q5 High-Fidelity DNA polymerase (NEB, M0494L) by following the program: 72°C for 5min, 98°C for 30 s; 5 cycles of 98°C for 10 s and 72°C for 90 s, 72°C for 2 min. The number of additional cycles needed for the PCR reaction was determined by qPCR - the number of qPCR cycles needed to reach 1/3 of the saturated signal was used as the additional PCR cycles. PCR products were purified using AMPure XP beads (Beckman Coulter; 0.7x–1.0x ratios depending on fragment size). The size distribution and concentration of the resulting libraries were assessed using an Agilent TapeStation and libraries were sequenced on an Illumina NextSeq or NovaSeq platform.

To perform ECHOS on tissue slices, tissue samples were cryo-sectioned at a thickness of 10 µm using a Leica CM1950 cryostat. Prior to mounting, 35-mm MatTek glass-bottom dishes (No. 1.5 coverslip, 14-mm diameter) were pre-coated with 0.1% poly-L-lysine (Sigma P4707) for 20 min and allowed air-dry completely. Freshly cut tissue slices were then directly transferred onto the poly-L-lysine–coated glass surface and were processed using the same protocol applied to cultured cells with minor modifications. For tissues slices, a higher concentration of loaded nb-Tn5 (1:12.5 dilution) was used. For tagmentation reactions, tissue sections were incubated for 3 hours at 37 °C.

### The ECHOS+ workflow

In ECHOS+, a photo-caged adaptor containing a partial T7 promoter (ECHOS_T7_75) was annealed with ME. nb-Tn5 loaded with the annealed adaptors were then used to tagment the genome in situ. Following photo-uncaging and DNA purification, ligation was performed to reconstitute the full length T7 promoter using the remaining T7 sequence (ECHOS_73) and a splint oligo (ECHOS_72). Following gap-filling, IVT was performed using the HiScribe T7 Quick kit (NEB, E2040L) at 37 °C for 14–16 h. The resulting RNA was purified using RNAclean XP beads (Beckman Coulter; 1.8× ratio) and eluted in 10 µL nuclease-free water. Purified RNA was reverse transcribed using random hexamer and Maxima H-Minus reverse transcriptase (Thermo Fisher, EP0753) at 50 °C for 30–60 min, followed by second-strand synthesis (SSS) using NEBNext 2× master mix (NEB, M0541L) and SSS primer (ECHOS_T7split_SSS) to generate double-stranded cDNA. The cDNA was used to generate sequencing libraries using an Illumina Nextera XT Library Preparation kit (FC-131-1096).

### CUT&Tag

The HeLa cell CUT&Tag experiment was performed using the CUT&Tag-IT Core Assay Kit (Active Motif, Catalog #53176) and following the manufacturer’s protocol. ∼100,000 HeLa cells were used as input for one reaction. The same primary antibodies as those used in the ECHOS experiments were applied to ensure consistency across platforms.

### Spatial transcriptomics using PHOTON

Human ectocervical epithelium compartment-specific gene expression profiles were generated using a spatial transcriptomics technology called PHOTON^37^. Briefly, tissue slices in a glass-bottom dish were fixed and permeabilized. Samples were stained with anti-Ki67 to aid the visualization of the luminal and basal compartment. In situ RT was performed overnight in the dark at 37 °C using photo-uncaged RT primers. The same photo-uncaging procedure as ECHOS was followed to photo-uncage the luminal and basal compartment, respectively. Afterwards, samples were digested and nucleic acids were then extracted. Annealed ligation adapters were ligated to the purified nucleic acids followed by Streptavidin bead pulldown. A second round of RT with the template switch oligos was performed followed by a PCR reaction. Purified PCR products were subject to sequencing library preparation using the Nextera XT Library Prep Kit.

### RNA FISH

For the human ectocervical samples, hybridization chain reaction (Molecular Instruments, HCR v3 kit) was performed by following the manufacturer’s protocol. Briefly, frozen OCT-embedded ectocervical tissue blocks were cryo-sectioned at 10 µm thickness and mounted directly onto positive-charged glass slides. Tissue sections were fixed in 4% PFA for 10 min, washed in PBS, and permeabilized in 70% ethanol at-22°C overnight. Sections were then pre-hybridized in HCR Hybridization Buffer for 30 min at 37 °C, followed by overnight hybridization with gene-speicific probe sets (4 nM) at 37 °C. The next day, sections were washed in pre-warmed Probe Wash Buffer (3–4 x 15 min, 37°C) and 5× SSCT. Fluorescent hairpins (h1/h2, snap-cooled) were applied in Amplification Buffer for 1–3 h at room temperature in the dark. After washing in 5x SSCT, nuclei were counterstained with DAPI and imaged on a Nikon A1R confocal microscope.

To simultaneously visualize XIST and H3K27me3 in the primary human skin fibroblasts, cells were plated onto Matrigel-coated glass-bottom plates, fixed in 4% PFA for 10 min, and permeabilized in 70% ethanol at –20 °C overnight. After rehydration in PBS, samples were blocked and incubated with anti-H3K27me3 primary antibody (CST 9733, 1:200) overnight at 4 °C, followed by an HCR-compatible secondary antibody for 1 h at room temperature. Cells were post-fixed in 4% PFA for 10 min. *XIST* RNA was detected following the same HCR v3 protocol as described above.

### ECHOS and ECHOS+ data analyses

1) Sequencing read processing and alignment

Sequencing reads were processed following similar pipelines established for ATAC-seq and CUT&Tag data^7,44^. Briefly, paired-end reads were adapter-trimmed using BBDuk (BBMap suite) and aligned using Bowtie2^45^ (v2.4.2) to the reference genome. Data generated from the HeLa cells were aligned to hg19. Data from the human skin fibroblasts and human ectocervical epithelium were aligned to hg38. MN datasets were aligned to the T2T-CHM13 genome. And the mouse cerebral cortex data were mapped to mm10. Bowtie2 was run in paired-end local alignment mode with parameters optimized for CUT&Tag-style tagmented fragments (maximum fragment length 2,000 bp). Aligned reads were sorted by genomic coordinate using Samtools^46^, and low-quality alignments (MAPQ ≤30) were excluded from downstream analysis. Unplaced or unlocalized contigs and mitochondrial DNA were removed where appropriate. PCR duplicates were identified and removed using Picard^47^ (MarkDuplicates), which was also used to estimate library complexity. Processed BAM files were used for all downstream analyses, including generation of normalized signal tracks, peak calling, and domain-level quantification.

2) Peak calling and domain identification

Peak calling was performed using MACS2^48^ (v2.1.2) with parameters matched to the expected architecture of each histone modification. Promoter-associated marks such as H3K27ac and H3K4me3 were called in narrow-peak mode, whereas broad histone domains such as H3K27me3 were identified using broad peak–calling settings. The resulting peak sets (narrowPeak or broadPeak) were used for feature quantification, heatmap generation, and differential analyses. For most ECHOS/ECHOS+ libraries—including those generated from the whole cells, the Barr bodies, and the tissue slices—peak calls were generated using MACS2 settiings consistent with CUT&Tag data analysis. MN analyses were performed differently because H3K27ac enrichment in MN manifests as broad, domain-like signals rather than discrete peaks. In this context, discrete peak calling is not biologically appropriate. Instead, MN analyses relied on normalized coverage tracks (CPM or RPGC) and bin-based domain quantification to capture broad H3K27ac patterns along the Y chromosome.

3) SNR quantification

Approximately 5000 cells were plated in each well of a 96-well glass-bottom plate. In situ tagmentation was performed following the ECHOS or ECHOS+ workflow. An incremental percentage of cells were photo-uncaged across the wells, starting from 0% (photo-caged control) to 10% (ECHOS) or 20% (ECHOS+). Sequencing libraries were then prepared for each well and sequenced. Reads were aligned and PCR duplicates were removed. The number of unique genomic fragments was used as a quantitative measure of library yield. We defined the noise fragments as the unique fragments in the photo-caged control library (N_noise_), and the signal fragments as the residual fragments in each photo-uncaged library after subtracting the noise (N_uncaged_ – N_noise_). The SNR was then calculated as N_uncaged_ – N_noise_ **/** N_noise_.

4) Differential peak analysis

Differential peak enrichment analysis was evaluated using a unified MACS2–HOMER–DESeq2 workflow. Peaks were first called using MACS2 for each replicate and merged to generate a union peak set for each group. Read coverage over the identified peaks were quantified using HOMER^49^ (annotatePeaks.pl), and differential analysis was performed using DESeq2^50^ via HOMER’s getDiffExpression.pl wrapper. DESeq2-derived log2 fold-changes and FDR-adjusted p-values were used to define differential peaks and to generate volcano plots, MA plots, and heatmaps for downstream interpretation.

5) MN hotspot analysis

MN–specific H3K27ac enrichment was quantified using normalized continuous signals, rather than discrete peak calls. SES-normalized log_₂_(MN/Control) BigWig tracks were generated using bamCompare (deeptools^51^) and were used for all downstream analyses. To detect chromatin domains with elevated H3K27ac signals in MN, the Y chromosome was divided into 5-kbp bins, and the SES-normalized signals were extracted from each MN and the corresponding control dataset for each bin. To quantify MN-specific gain relative to the nuclear background, a differential score was computed as: ΔR = R_MN_ - R_Ctrl_, where R_MN_ and R_Ctrl_ denote the MN and control nuclear H3K27ac signals, respectively. Bins exceeding an arbitrary threshold for both the absolute signal level (>0.6) and the differential score (ΔR > 0.6) were classified as hotspots. Adjacent bins were merged into continuous domains using a gap ≤10-kbp rule and smoothed 50-kbp profiles were used for visualization.

6) Xi H3K27me3 profiling

SES-normalized log_₂_(Barr body/Control) bigwig tracks were generated for the young and aging samples using bamCompare. To assess regional H3K27me3 patterns across the Xi, chrX was divided into non-overlapping 1-Mb windows, and the mean normalized signal within each window was computed from the bigwig tracks. Window-level young and aging signals were plotted along the chromosomal coordinates, and window-wise signal differences (Aging − Young) and log_₂_ ratios (Aging/Young) were calculated for comparative visualization.

### PHOTON data processing

The spatial transcriptomics data were preprocessed using the PHOTON analysis pipeline^37^. Briefly, trimmed sequencing reads were aligned to the human reference genome (hg38) using HISAT2^52^, quantified with featureCounts^53^, and analyzed for differential expression using DESeq2. Differentially expressed genes (adjusted p value < 0.05) were used for integrative comparison with ectocervical epithelial compartment-specific ECHOS+ epigenetic profiles.

## Data availability

The raw sequencing data supporting the findings of this study are available in the NCBI BioProject database with the BioProject ID # PRJNA1373458. The accession # of the public HeLa cell LaminB1 ChIP-seq data is GSE57149. The accession # of the public mouse cerebral cortex H3K27ac ChIP-seq data GSE231221. The accession # of the public HeLa cell H3K27ac and H3K4me3 ChIP-seq data is GSE29611.

## Code availability

Custom code is available at https://github.com/HaiqiChenLab/ECHOS.

## Acknowledgements

H.C. acknowledges support from the Cecil H. and Ida Green Center for Reproductive Biology Sciences Endowment and NIH/NHGRI (R01HG013358). M.M. acknowledges support by the NIH/NICHD (R01HD110147). E.J.G. acknowledges support from CPRIT (RR210077-Grow) and NIH/NIGMS (R35GM159821). We thank Dr. Maria Florian-Rodriguez for providing human cervical tissues through the Human Tissue, Biological Fluids and Cell Culture Laboratory Core in the Department of Obstetrics and Gynecology at UT Southwestern Medical Center. We also thank members of the Peter Ly Lab at UT Southwestern Medical Center for the help with the micronucleus experiments.

## Author contributions

H.C. conceived and supervised the project. Q.C. performed experiments and analyzed the data. Q.X., Y.U., S.R., M.S., and X.Z. assisted with the experiments. M.M. and E.J.G. provided consultation. H.C. and Q.C. wrote the manuscript with input from all authors.

---

**Figure S1.**
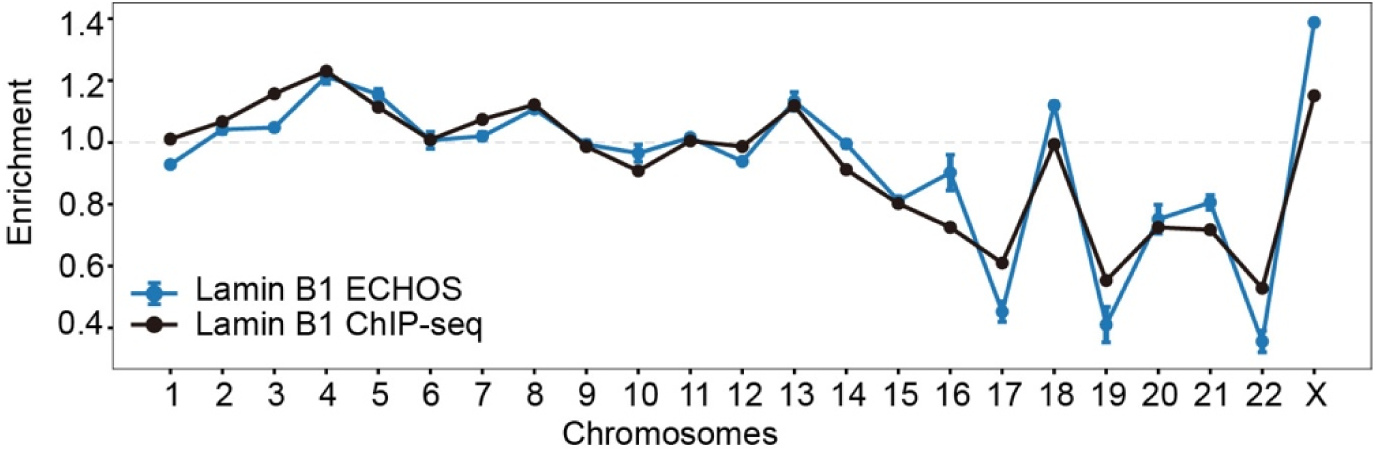
ECHOS recapitulates the genome-wide enrichment profile of Lamin-associated chromosomes. ECHOS-generated nuclear periphery enrichment profile compared with a published Lamin B1 ChIP–seq dataset at the chromosome level.

**Figure S2.**
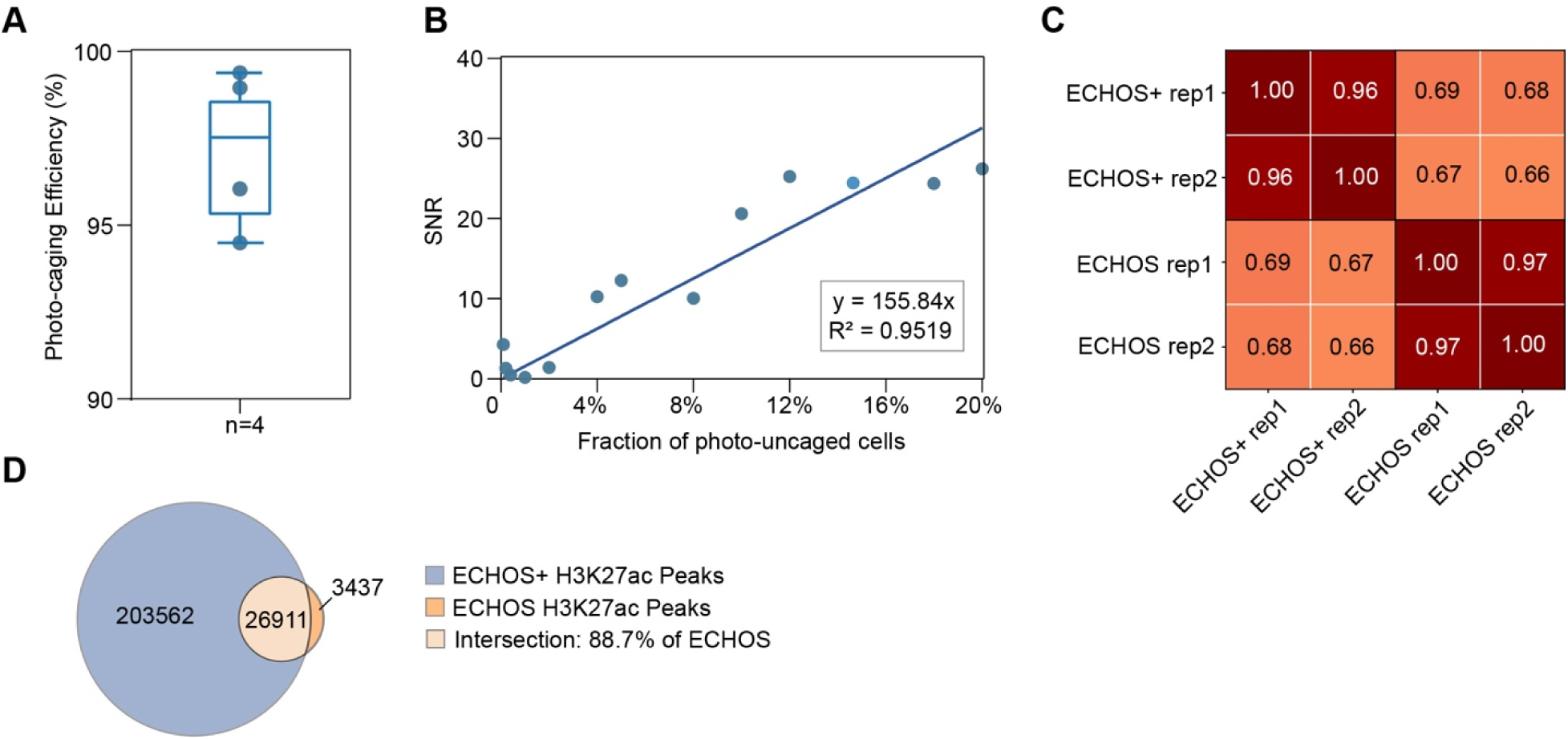
ECHOS+ enhances library complexity while preserving specificity. **A** Photo-caging efficiency for ECHOS+ across 4 biological replicates. **B** SNR as a function of the fraction of cells photo-uncaged using ECHOS+. Each point represents an independently prepared library, with linear regression shown (slope = 155.84; R² = 0.9519). **C** Pairwise Pearson correlation matrix computed from genome-wide 10-kbp binned coverage for H3K27ac datasets generated by ECHOS and ECHOS+, respectively. **D** Venn diagram showing the overlap between ECHOS+ and ECHOS H3K27ac peaks.

**Figure S3.**
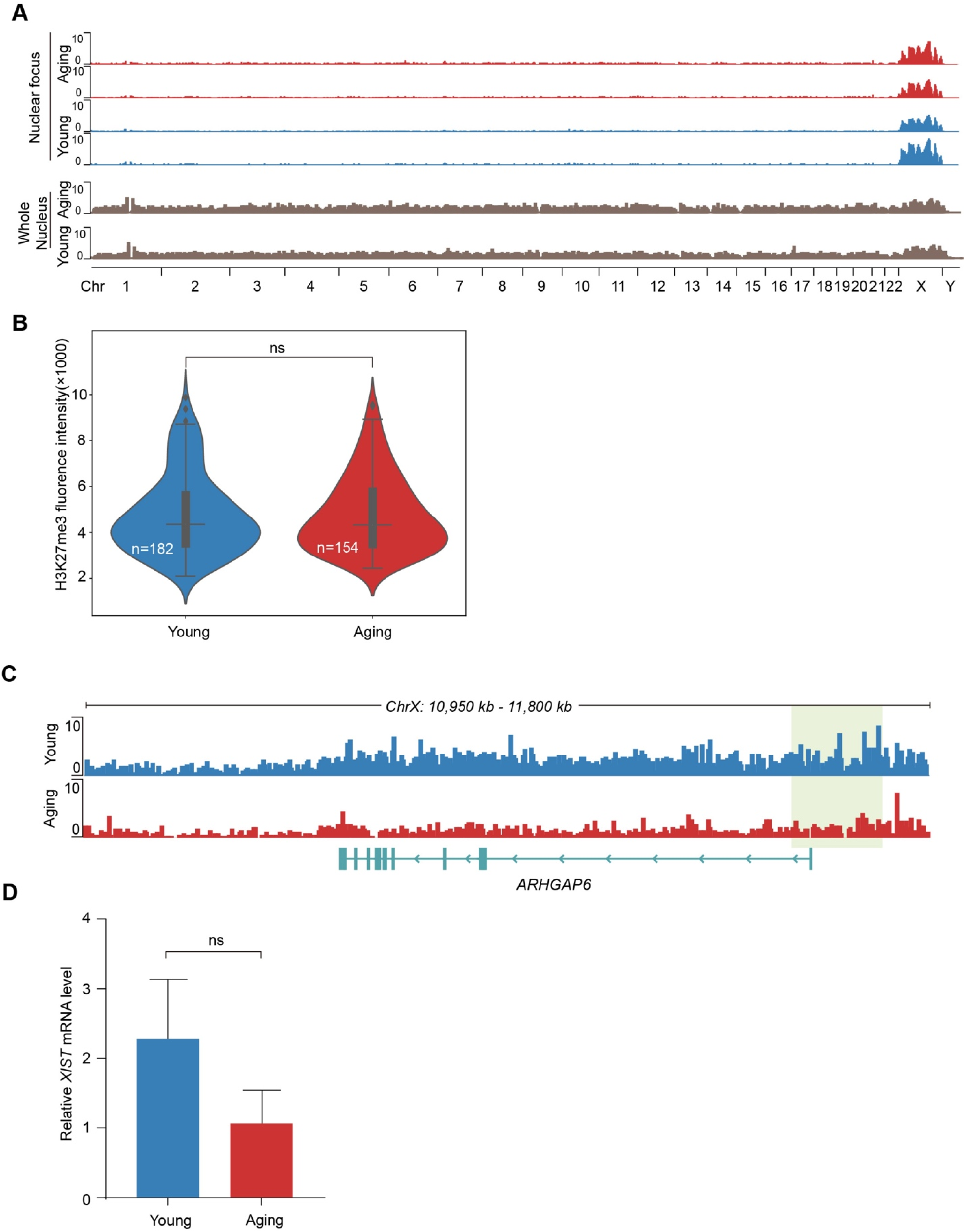
ECHOS+ captures the H3K27me3 profile of the Xi **A** Genome-wide normalized H3K27me3 coverage tracks (50-bp bins) of the nuclear focus and the primary nucleus from young and aging female skin fibroblasts. **B** H3K27me3 fluorescence intensity within the Barr body of young (n=182) and aging (n=154) fibroblasts (Mann–Whitney U test, p=0.840). ns, not significant. **C** H3K27me3 genome browser views of the ARHGAP6 locus in both the young and aging group. **D** qPCR result of the XIST expression level in young and aging cells. ns, not significant.

