## Supplementary material for "ECHOS enables spatial epigenome profiling at subcellular resolution": Table S1

**Table S1. Oligonucleotides used in ECHOS and ECHOS+**

**Adaptor and ligation oligos for ECHOS**

| **Name** | **Sequence (5’→3’)** | **Chemical modifications** | **Function / Role** | **Used in** | **Notes** |
| --- | --- | --- | --- | --- | --- |
| **ECHOS_17** | /5Alex546N//iSpPC/TCGTCGGCAGCGTCAGATGTGTATAAGAGACAG | 5’ Alexa546, internal iSpPC | Forward (P5) adapter | ECHOS | Photocleavable |
| **ECHOS_21** | /5Alex546N//iSpPC/GTCTCGTGGGCTCGGAGATGTGTATAAGAGACAG | 5’ Alexa546, internal iSpPC | Reverse (P7) adapter | ECHOS | Photocleavable |
| **ECHOS_37** | /5phos/CTGTCTCTTATAC/3ddC/ | 5’ phosphate, 3’ dideoxyC | Mosaic-end (ME) | ECHOS, ECHOS+ | – |
| **ECHOS_22** | AATGATACGGCGACCACCGAGATCTACACCTTCTCTAT | – | P5 oligo | ECHOS | – |
| **ECHOS_24** | TGCCGACGAATAGAGAGGTTGTAG/3ddC/ | 3’ dideoxyC | P5 splint | ECHOS | ligation splint oligo |
| **ECHOS_39** | CCCACGAGACAGACAGTCAGTCG/3ddC/ | 3’ dideoxyC | P7 splint | ECHOS | ligation splint oligo |
| **ECHOS_40** | CAAGCAGAAGACGGCATACGAGAT | – | P7 PCR primer | ECHOS | – |
| **ECHOS_57** | AATGATACGGCGACCACCGAGAT | – | P5 PCR primer | ECHOS | – |

**Index sequence for ECHOS**

| **Name (ECHOS_IDX)** | **Sequence (5’→3’)** | **Index (adapter)** | **Index (sample sheet)** | **Function** |
| --- | --- | --- | --- | --- |
| ECHOS_IDX_S501 | CAAGCAGAAGACGGCATACGAGATTAGATCGCCGACTGACTCT | TAGATCGC | GCGATCTA | i7 index |
| ECHOS_IDX_S502 | CAAGCAGAAGACGGCATACGAGATCTCTCTATCGACTGACTCT | CTCTCTAT | ATAGAGAG | i7 index |
| ECHOS_IDX_S503 | CAAGCAGAAGACGGCATACGAGATTATCCTCTCGACTGACTCT | TATCCTCT | AGAGGATA | i7 index |
| ECHOS_IDX_S506 | CAAGCAGAAGACGGCATACGAGATACTGCATACGACTGACTCT | ACTGCATA | TATGCAGT | i7 index |
| ECHOS_IDX_S507 | CAAGCAGAAGACGGCATACGAGATAAGGAGTACGACTGACTCT | AAGGAGTA | TACTCCTT | i7 index |
| ECHOS_IDX_S508 | CAAGCAGAAGACGGCATACGAGATCTAAGCCTCGACTGACTCT | CTAAGCCT | AGGCTTAG | i7 index |
| ECHOS_IDX_N701 | CAAGCAGAAGACGGCATACGAGATGCGTAAGACGACTGACTCT | GCGTAAGA | TCTTACGC | i7 index |
| ECHOS_IDX_N702 | CAAGCAGAAGACGGCATACGAGATTCGCCTTACGACTGACTCT | TCGCCTTA | TAAGGCGA | i7 index |
| ECHOS_IDX_N703 | CAAGCAGAAGACGGCATACGAGATCTAGTACGCGACTGACTCT | CTAGTACG | CGTACTAG | i7 index |
| ECHOS_IDX_N704 | CAAGCAGAAGACGGCATACGAGATTTCTGCCTCGACTGACTCT | TTCTGCCT | AGGCAGAA | i7 index |
| ECHOS_IDX_N705 | CAAGCAGAAGACGGCATACGAGATGCTCAGGACGACTGACTCT | GCTCAGGA | TCCTGAGC | i7 index |
| ECHOS_IDX_N706 | CAAGCAGAAGACGGCATACGAGATAGGAGTCCCGACTGACTCT | AGGAGTCC | GGACTCCT | i7 index |
| ECHOS_IDX_N707 | CAAGCAGAAGACGGCATACGAGATCATGCCTACGACTGACTCT | CATGCCTA | TAGGCATG | i7 index |
| ECHOS_IDX_N708 | CAAGCAGAAGACGGCATACGAGATGTAGAGAGCGACTGACTCT | GTAGAGAG | CTCTCTAC | i7 index |
| ECHOS_IDX_N709 | CAAGCAGAAGACGGCATACGAGATCCTCTCTGCGACTGACTCT | CCTCTCTG | CAGAGAGG | i7 index |
| ECHOS_IDX_N710 | CAAGCAGAAGACGGCATACGAGATAGCGTAGCCGACTGACTCT | AGCGTAGC | GCTACGCT | i7 index |
| ECHOS_IDX_N711 | CAAGCAGAAGACGGCATACGAGATCAGCCTCGCGACTGACTCT | CAGCCTCG | CGAGGCTG | i7 index |
| ECHOS_IDX_N712 | CAAGCAGAAGACGGCATACGAGATTGCCTCTTCGACTGACTCT | TGCCTCTT | AAGAGGCA | i7 index |

**Adaptor and ligation oligos for ECHOS+**

| **Name** | **Sequence (5’→3’)** | **Chemical modifications** | **Function / Role** |
| --- | --- | --- | --- |
| **ECHOS_T7_75** | /5Cy3//iSpPC/CGAC/iNPOM-dT/CACTATAGGGTCGTCGGCAGCGTCAGATGTGTATAAGAGACAG | Cy3, iSpPC, iNPOM-dT | Partial T7 promoter-containing adapter |
| **ECHOS_37** | /5phos/CTGTCTCTTATAC/3ddC/ | 5phos, 3ddC | ME |
| **ECHOS_73** | GAATTTATAA | – | The remaining T7 promoter sequence |
| **ECHOS_72** | GACGCCACTATAGGATGTGTITATAAATTC/3ddC/ | 3’ dideoxyC | The splint to ligate the remaining T7 promoter sequence to the adaptor |
| **ECHOS_7split_SSS** | GTCGTGGCAGCGTC | – | Second-strand synthesis primer |
